# Customizable Live-Cell Imaging Chambers for Multimodal and Multiplex Fluorescence Microscopy

**DOI:** 10.1101/2020.02.19.955971

**Authors:** Adam Tepperman, David Jiao Zheng, Maria Abou Taka, Angela Vrieze, Austin Le Lam, Bryan Heit

## Abstract

Using multiple imaging modalities while performing independent experiments in parallel can greatly enhance the throughput of microscopy-based research, but requires provision of appropriate experimental conditions in a format that meets the microscopy’s optical requirements. Although customized imaging chambers can meet these challenges, the difficulty of manufacturing custom chambers and the relatively high cost and design inflexibility of commercial chambers has limited the adoption of this approach. Herein, we demonstrate the use of 3D printing to produce inexpensive, customized live-cell imaging chambers that are compatible with a range of imaging modalities including super-resolution microscopy. In this approach, biocompatible plastics are used to print imaging chambers designed to meet the specific needs of an experiment, followed by adhesion of the printed chamber to a glass coverslip, producing a chamber that is impermeant to liquids and which supports the growth and imaging of cells over multiple days. This approach can also be used to produce moulds for casting PDMS microfluidic devices. The utility of these chambers is demonstrated using designs for multiplex microscopy, imaging under shear, chemotaxis, and general cellular imaging. Together, this approach represents an inexpensive yet highly customizable approach to produce imaging chambers that are compatible with modern microscopy techniques.

## Introduction

Technological advances have enabled a broad array of live-cell imaging approaches to investigate cellular processes, many of which require optically demanding imaging modalities to achieve the necessary resolution and sensitivity ^1^. Achieving these optical conditions usually requires the use of high numerical aperture oil-immersion objective lenses, which are often designed to work with coverslips of a very specific thickness and refractive index – usually a #1.5 (0.17 mm) coverslip of η = 1.515, with deviation from these parameters compromising the resolution of the resulting images ^2,3^. These imaging modalities then need to be employed in an experimental system capable of providing the necessary experimental conditions, for example, shear flow ^4^, chemoattractant gradient generation ^5^, introduction of stimuli ^6^, electrophysiology electrodes ^7^, and multiple wells for multiplex or multimodal imaging ^8^. While commercial imaging chambers of varying design are available, compromises are often made between identifying a chamber that meets the physical requirements of an experiment versus meeting the optical requirements of the microscopy. As one example, some commercial chambers use plastic coverslips which do not exactly match the optical characteristics of #1.5 glass coverslips and are incompatible with some immersion oils, leading to image degradation and precluding their use with some forms of microscopy.

Many research groups have overcome these limitations by producing imaging chambers in-house. Historically, two manufacturing approaches have been used. Milling of aluminum or plastic blocks to produce chambers is a common approach, but is limited in its ability to produce small features, and it is extremely difficult to produce interior features ^9^. Moreover, this approach can be expensive, especially for single-use devices as making these devices requires access to a CNC milling machine and costly machining blanks made of biocompatible materials. A second approach is the use of moulds to cast chambers out of polydimethylsiloxane (PDMS), a biocompatible silicone polymer ^10^. PDMS molds can be produced by milling or by photolithography, with the latter offering much higher resolution and a better ability to generate complex structural features ^11^. While most PDMS chambers are limited to features formed between voids in the cast and the underlying coverslip, devices with complex internal features can be produced by layering multiple PDMS casts ^12^. However, photolithography is a specialized technique that can be cost-prohibitive, and chamber assembly requires an oxygen plasma generator to covalently bond the PDMS chamber to a glass coverslip ^13^. Thus, while custom-designing and manufacturing chambers is an ideal solution for many microscopy experiments, the adoption of this approach has been limited due to the relatively high cost and expertise required to create these chambers.

The recent proliferation of consumer-oriented fused deposition modeling (FDM) 3D printers offers the opportunity to easily design and manufacture customized imaging chambers in a cost-effective manner. FDM is an additive manufacturing process in which a thermoplastic filament is heated above its glass transition temperature and extruded as a thin layer onto the build surface ^14^. Printed products are built via the sequential addition of layers of materials, allowing for complex three-dimensional objects to be built, including objects with complex internal structures. Consumer-oriented FDM printers generally have a maximum z-resolution (e.g. layer height) of mm to 0.10 mm, with the lateral resolution determined by the diameter of the print nozzle – typically 0.2 mm to 0.4 mm. This resolution is intermediary between that of milling and photolithography, at a price-point well below either of these approaches. Several free and inexpensive software packages are available for designing 3D printed objects, and printing can be performed with biocompatible and food-grade materials such as polylactic acid (PLA), polycarbonate (PC) and nylon ^15^. Indeed, other groups have used this approach to print imaging chambers for culturing and stimulating neurons ^7^, general cell culture ^16^, zebrafish imaging ^8^ and for processing and imaging clarified tissues ^17^. However, many of these approaches image the sample via a water-dipping lens or suspend the sample in a water-based gel, thus limiting their use for high-resolution and super-resolution microscopy.

Herein, we present and validate a general approach for creating 3D printed imaging chambers which are mounted on conventional glass coverslips. These are fully customizable chambers that can contain internal structures, are suitable for live-cell imaging, can be designed for multiplex and multimodal imaging, and enable high-resolution and super-resolution microscopy without any loss in resolution or precision.

## Results

### Identification of Suitable Chamber Materials

Many plastics can be used for FDM printing, but of these, only PLA has consistently been shown to be cell compatible ^7,16,18^. To test the cell culture suitability of commonly used FDM plastics, we cultured HeLa and J774.2 cells with PLA, nylon, PC and glycol-modified polyethylene terephthalate (PETG) for 5 days and, using a crystal violet cytotoxicity assay, found no evidence of overt toxicity by any of the plastics (**Figure 1A**). While this experiment found no signs of cytotoxicity, it is possible that these plastics may cause more subtle cellular effects. To assess this possibility, we analyzed actin and plasma membrane morphology after 48 hours of culture with these plastics, using adjacency statistics as a sensitive measure of morphological changes resulting from cell stress or other effects of the plastics on cell morphology ^19^. Adjacency statistics provides an unbiased measure of the spatial patterning and distribution of fluorescent markers by, for each pixel above an intensity threshold, quantifying the number of neighbouring pixels also above the threshold. This provides a 9-dimensional morphological readout per fluorescent channel (e.g. fraction of pixels with 0, 1, 2…8 above-threshold neighbours), which when calculated across three thresholds, provides a 27-dimensional measure of the morphology of a single fluorescent marker. The resulting 54-dimensional (actin + plasma membrane) morphology of the cells in our assay was reduced to 2-dimensions using principal component analysis, and was readily able to identify pre-apoptotic morphological changes induced by the pan-kinase inhibitor staurosporine in HeLa cells (**Figure 1B-C**). As previously reported, PLA had no apparent effect on cells cultured in its presence, and much to our surprise, the other tested plastics displayed a similar degree of cell compatibility (**Figure 1D-H**). Given the performance of PLA, and its successful use by other groups, we used this plastic for all future experiments. While we have successfully made imaging chambers from multiple brands of PLA from different manufactures (data not shown), for consistency all experiments presented in this study were performed using a single brand of PLA filament.

**Figure 1:**
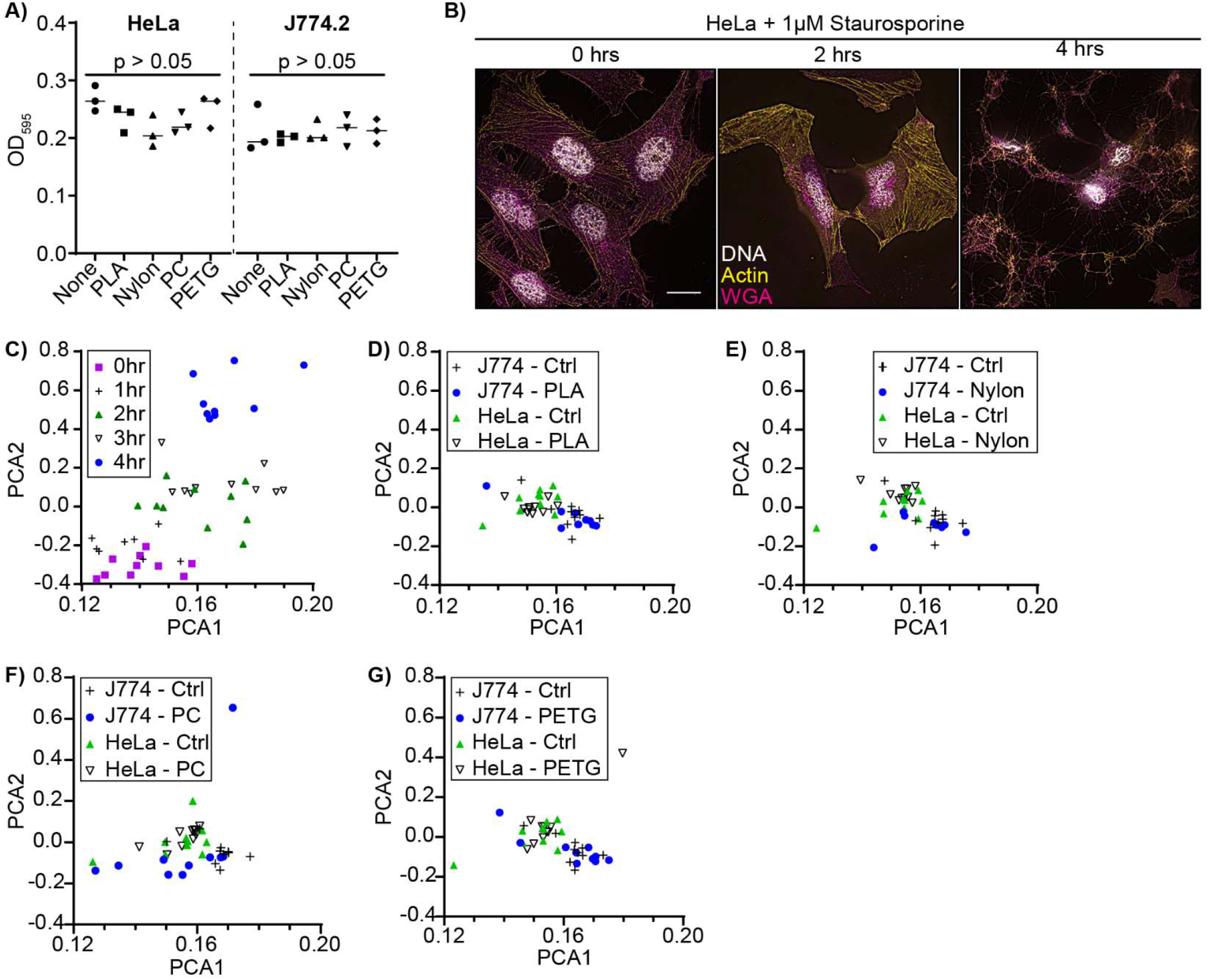
Determination of FDM Plastic Cytotoxicity. HeLa and J774 cells were assessed for cytotoxicity after culture with various plastics. **A)** Crystal violet viability assay after 5 days culture with the various plastics. **B-C)** representative SRRF images (B) and quantification of morphological changes with adjacency statistics (C) of HeLa cells over time following treatment with 1 μM staurosporine (STS). Cells are stained for DNA with Hoechst, actin with phalloidin-AlexaFluor-555, and the plasma membrane with wheat-germ agglutinin-AlexaFluor-647 (WGA). Scale bar is 10 μm. **D-G)** Morphological changes of J774.2 and HeLa cells quantified using adjacency statistics following 48 hr co-culture with PLA (D), nylon (E), PC (F), and PETG (G). Data is plotted as individual measurements from repeat experiments (A) or as principal components of the aggregate data collected from individual images collected over 3 independent experiments (C-G). p-values are calculated using a one-way ANOVA with Tukey test.

### Chamber Stability and Precession

Many live-cell experiments require water-tight or air-tight chambers to allow for fluid flow, experiments under pressure/vacuum, or control over atmospheric conditions. Running contrary to these needs are two issues: firstly, FDM printing produces weaker structures than does injection molding with the same plastic ^20^. Secondly, unlike PDMS, FDM printed parts cannot be covalently linked to glass as they are destroyed by oxygen plasma treatment, and our attempts to chemically crosslink prints to functionalized glass were not successful (data not shown) ^21^. Cyanoacrylate glues are often used to assembled FDM printed parts, but cells underwent rapid cell death when placed in chambers where the coverslip was attached with this adhesive (**Figure 2A**). As such PDMS was the only available biocompatible adhesive for chamber assembly ^22^. To test the strength of FDM prints and PDMS bonding an ^18^ mm square coverslip was attached with PDMS over the 10 mm × 10 mm opening of a 3D printed chamber with a 0.5 cm^3^ internal volume, creating a chamber with an ~225 mm^2^ glued surface. This chamber was filled with water and then pressurized to 400 kPa in 50 kPa intervals (**Figure 2B**). The chambers failed between 280 kPa and 370 kPa due to cracking of the coverslip; prior to this point no evidence of water leakage through the chamber material or PDMS seal was observed. Next, the chambers were subjected to a vacuum and the chamber pressure monitored. A slow loss of vacuum was observed (~0.8 kPa/min, **Figure 2C**), and interestingly this was also observed in a chamber comprised entirely of PLA, suggesting that this loss was due to gas permeability of the PLA itself and not due to leakage through the PDMS seal.

**Figure 2:**
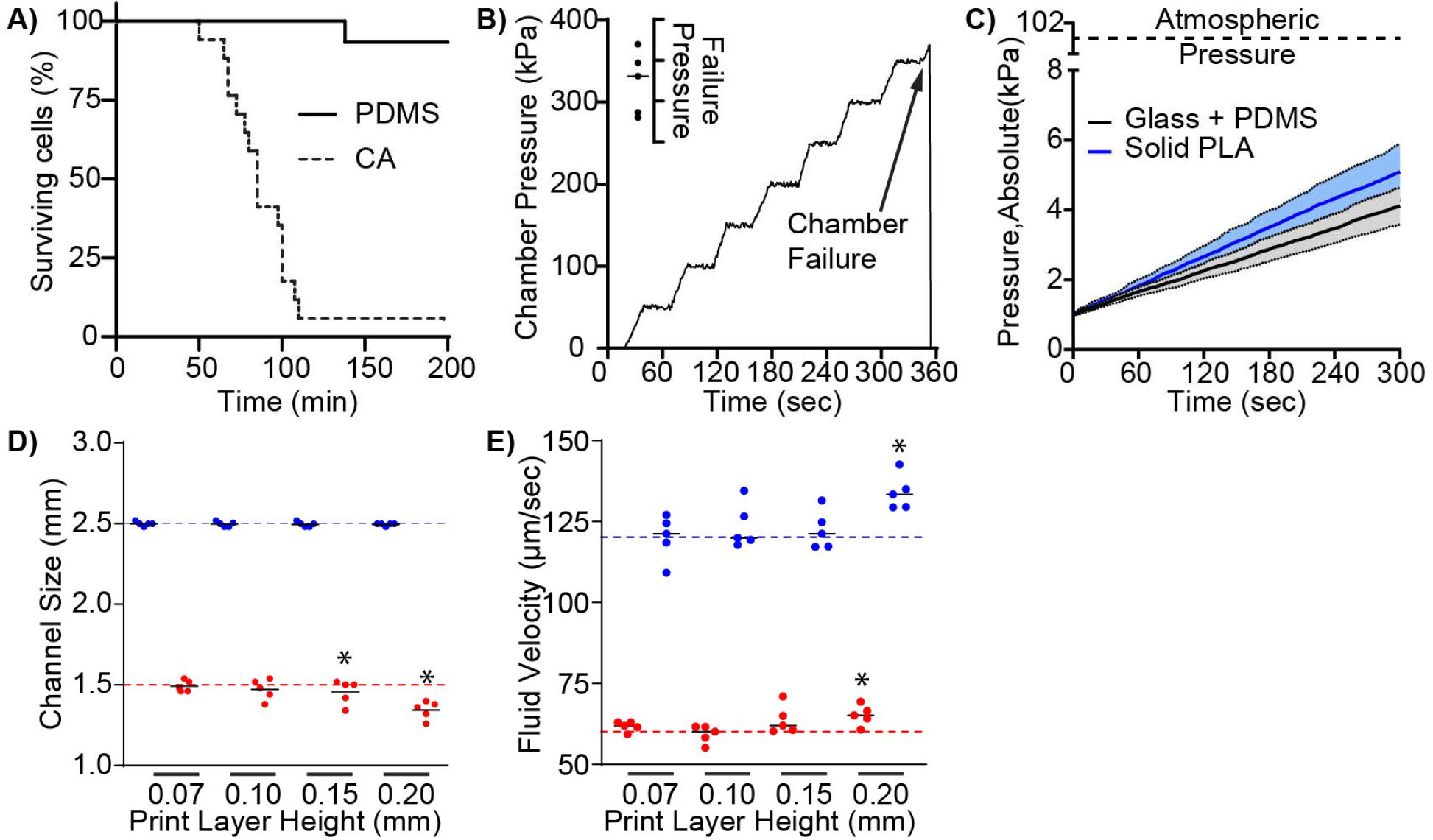
Characterization of Chamber Strength and PrintAccuracy. **A)** Survival of RAW264.7 macrophages during imaging in chambers assembled with cyanoacrylate (CA) or PDMS. n = 15. **B)** Representative pressurization trace of PLA chambers sealed with a #1.5 thickness coverslip and PDMS. Insert shows the failure pressure of 5 independent chambers. **C)** Air permeability of PLA chambers under vacuum. Chambers were sealed either with a coverslip and PDMS (Glass + PDMS) or were printed as an equal volume chamber completely enclosed in PLA (Solid PLA). Data is plotted as the mean chamber pressure ± 95% CI of 5 independent chambers/condition. **D-E)** Effect of print layer height on the print accuracy of a flow chamber with channels 2.5 mm in width and 1.5 mm in height. **D)** Measured channel width (blue) and height (red) compared to the design dimensions of the chamber (dotted lines). E) Effect of print layer height on the fluid flow velocity through the chamber. Chambers were perfused at two flow rates, which would produce theoretical flow velocities of 60 μm/s (red line) and 120 μm/s (blue line). Data is plotted as the mean flow velocity measured in 5 independent chambers with the horizontal lines indicating the mean of the 5 chambers, * p > 0.05 compared to PDMS (A) or 0.07 mm layer height at the same dimension or flow rate (D-E), Kaplan-Meier test (A) or ANOVA with Tukey correction (D-E).

Finally, accurate chamber sizing is required for some applications – e.g. flow chambers where specific shear rates are required. To assess the accuracy of FDM printed chamber construction, we prepared flow chambers bearing flow channels 16 mm in length, 2.5 mm in width and 1.5 mm in height which were printed using layer heights of 0.07 mm to 0.20 mm. Measurements of the chambers with a high-precision calliper demonstrated that small layer heights (<0.1 mm) produced chambers of the expected dimensions, while larger layer highest produced chambers with the expected width but with heights slightly lower than designed (**Figure 2D**). To determine the effect of print accuracy on chamber performance, we captured high-speed time-lapse micrographs of 5 μm silica beads perfused through the chambers at two flow rates that would producing fluid velocities of 60 μm/s and 120 μm/s in chambers whose dimensions did not deviate from the chamber design (**Figure 2E**). Smaller layer heights (e.g. finer resolution printing) produced chambers that consistently produced the expected fluid velocities, while larger print heights resulted in higher than predicted flow rates, consistent with the lower channel height in these chambers. Combined, these data demonstrate that, when printed with a small layer height, FDM printing produces durable chambers of predictable dimensions that are water-tight, but with a low degree of gas permeability. These characteristics make these chambers suitable for experiments conducted under physiological flow rates and pressures.

### High-Resolution and Super-Resolution Imaging

Next, we assessed the suitability of FDM printed imaging chambers for high-resolution and super-resolution applications. First, we performed super-resolution radial fluctuation (SRRF) microscopy on phalloidin-stained HeLa cells. Phalloidin-actin complexes are 10-12 nm in diameter ^23^, allowing us to estimate the resolution of our SRRF reconstructions based on the full-width at half-maximum (FWHM) of a line profile taken perpendicular to a labeled actin filament (**Figure 3A**). FDM chambers bearing #1.5 thickness coverslips produced SRRF reconstructions with higher resolution than commercial chambers bearing a plastic coverslip of similar thickness and refractive index (**Figure 3B**). Indeed, SRRF reconstructions with our FDM chambers exceeded the Airy resolution of our microscope (*r*_*Airy@580 nm*_ = 253 nm, *r*_*SRRF*_ = 231 nm), whereas the commercial chamber provided a resolution similar to that of a confocal acquisition (280 nm). Next, we used our single-particle tracking approach to assess the suitability of FDM chambers for Single Molecule Localization Microscopy (SMLM) methods. HeLa cells expressing HA-tagged CD93 were labeled at low density with anti-HA Fab fragments directly conjugated to Cy3. The diffusion of these molecules was then imaged at a high frame rate, and the resulting image series subjected to SMLM reconstruction using a mixed-model fitting approach ^24^. The first step of this process is to measure the point-spread function (PSF) of the microscope by mapping Gaussian curves to local maxima. Our FDM chambers produced measured PSFs closer to the theoretical PSF of our imaging system than did plastic-bottomed chambers (**Figure 3C**), which in turn, produced more precise protein localizations (19.83 nm vs. 29.27 nm, **Figure 3D**). Combined, these data demonstrate that FDM printed chambers are suitable for high-resolution and super-resolution microscopy approaches and have superior performance to plastic-bottomed chambers.

**Figure 3:**
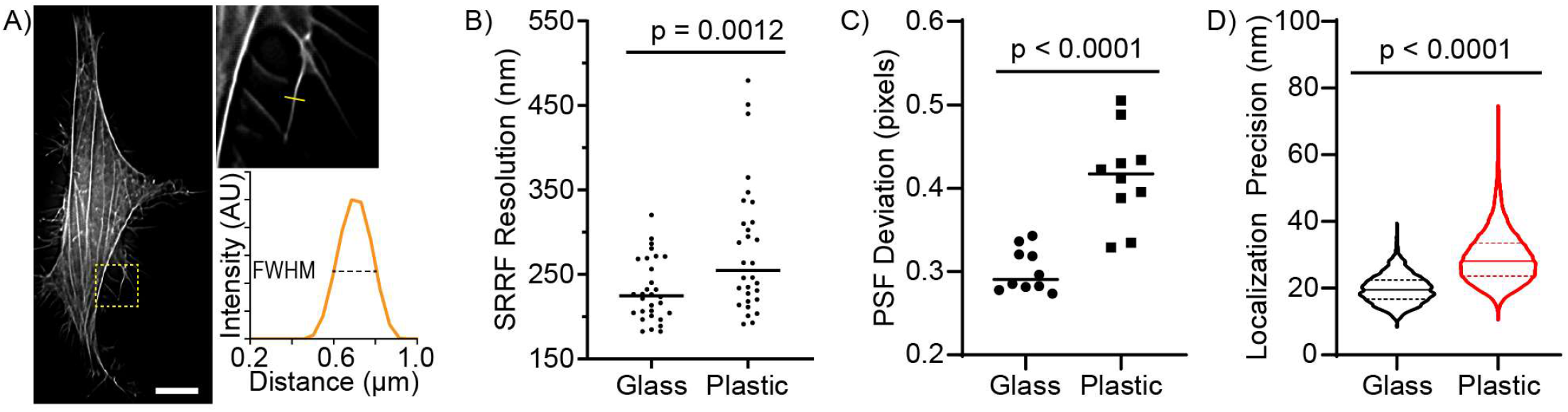
High-Resolution and Super-Resolution Microscopy in FDM Printed Chambers. The performance of glass-bottomed FDM printed chambers was compared to a commercial plastic-bottomed imaging chamber using SRRF and SMLM approaches. **A)** Estimation of SRRF image resolution using phalloidin-labeled actin in HeLa cells. A line profile is drawn perpendicular across a small actin fibril (yellow line shown in the expanded region of interest), with the image resolution estimated as the FWHM of the resulting intensity profile. Scale bar is 10 μm, expanded region of interest is 9.8 × 9.8 μm. **B)** Resolution of SRRF acquisitions on glass versus plastic coverslips. Cells on both types of coverslips were imaged with identical microscope settings and reconstructed using identical software settings. **C)** Deviation from theoretical of the PSF’s in SMLM acquisitions in imaging chambers with glass and plastic coverslips. **D)** Localization precision of individual fluorophores in an SMLM acquisition of cells on glass versus plastic coverslips. Data is representative of or quantifies (A-B) the ensemble of individual actin fibrils measured in 3 independent experiments, (C) the average PSF calculated from 10 SMLM acquisitions, or (D) the ensemble of these acquisitions across 3 independent experiments, quantifying a minimum of 100,000 detections per acquisition. Solid horizontal lines indicate the median, dotted horizontal lines indicate the 25^th^/75^th^ quartiles. p-values were calculated with a 2-tailed Mann-Whitney test.

### Multiplex & Multimodal Imaging of Phagocytosis

The ease with which custom-designed chambers can be created by FDM printing offers many opportunities to design imaging chambers matching the specific needs of an experiment. To illustrate this potential, we created an imaging chamber for multiplex and multimodal live-cell imaging of samples where small cell numbers are available (**Figure 4A**). This chamber features a 3 × 3 array of wells that fits onto an 18 mm × 18 mm coverslip, with each well completely separated from its neighbours. This allows for multiplex imaging of up to ^9^ separate experiments, using as few as 500 cells/well. Using this chamber and microscope acquisition software capable of scripting different acquisition parameters for each well, we simultaneously performed multiple assays of macrophage antimicrobial function, using a combination of white-light, fluorescence and ratiometric microscopy to illustrate these chambers multimodal capabilities.

**Figure 4:**
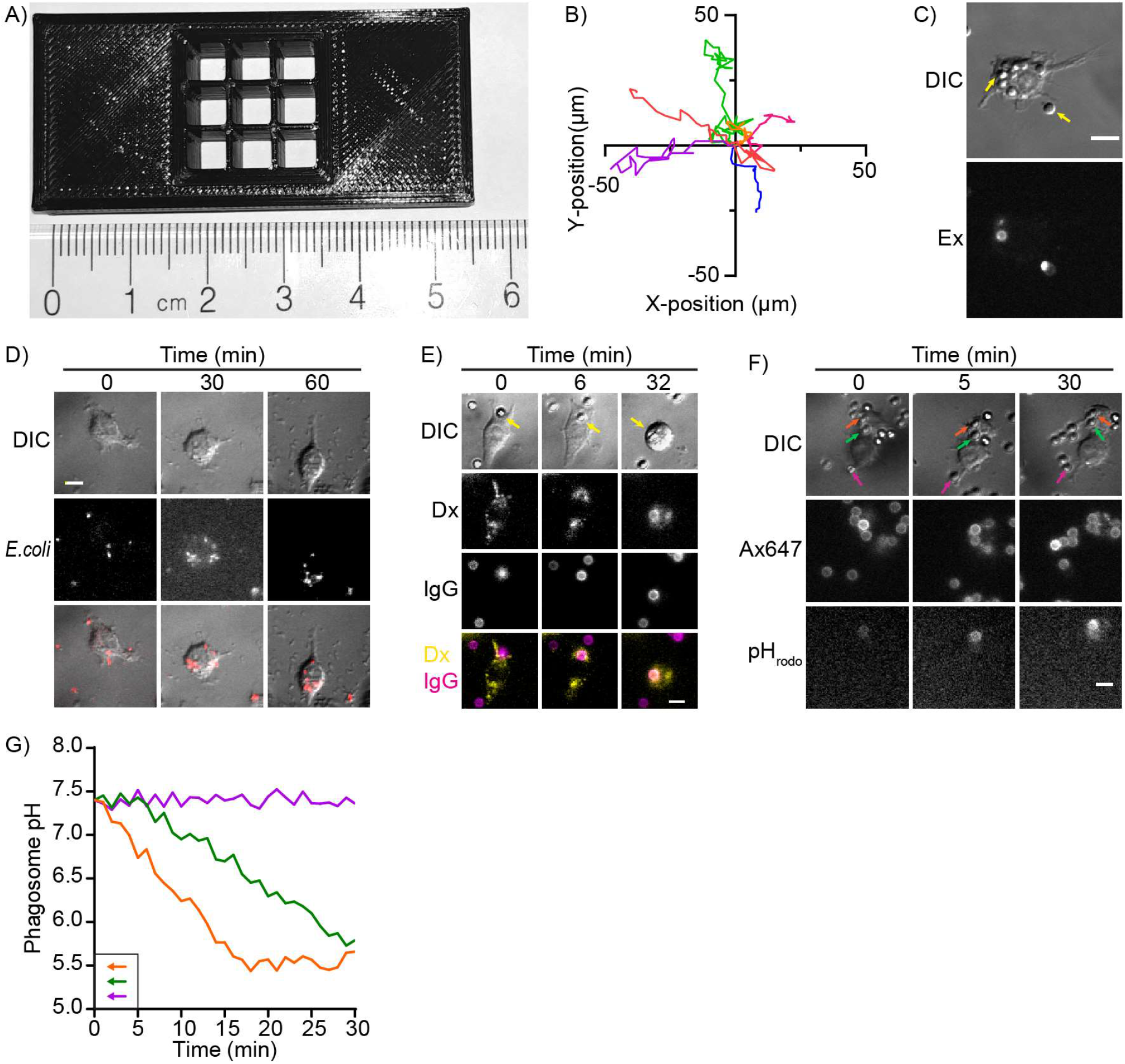
Multiplex & Multimodal Imaging of Macrophage Function using a Custom Printed Chamber. **A)** Design of a multiplex imaging chamber. J774.2 macrophages were placed into each well and multiple live-cell and end-point assays of macrophage function performed simultaneously over a period of 1 hr. **B)** Spontaneous migration of macrophages was quantified by DIC imaging and manual tracking of cell positions. **C)** An end-point phagocytosis assay was performed using IgG coated beads as pathogen mimics; non-phagocytosed beads were detected by staining with a fluorescent anti-IgG antibody at the end of the experiment (Ex & arrows). **D)** Time-lapse micrographs of a single macrophage phagocytosing and degrading fluorescently labeled *E. coli*. E) Quantification of lysosome-phagosome fusion using macrophages with fluorescent-dextran containing lysosomes (Dx) and IgG coated beads as pathogen mimics. Lysosome-phagosome fusion can be observed in phagosomes as an accumulation of dextran around the mimics (arrows). **F)** Ratiometric imaging of IgG-coated pathogen mimics labeled with a pH sensitive (pHrodo) and insensitive (Ax-647) fluorophores. Coloured arrows track two phagocytosed (green, orange) and one non-phagocytosed (purple) mimics. **G)** Acidification of the pathogen mimics tracked in panel F. Data is representative of 5 independent experiments, C-F: scale bars are 10 μm.

Macrophages actively patrol tissues ^25^, with this spontaneous migration quantified using a white light microscopy based cell-tracking assay (**Figure 4B**). Once a pathogen is encountered, the macrophage engulfs the pathogen through phagocytosis ^26^. The efficiency of phagocytosis was measured using an end-point assay and IgG-coated beads as pathogen mimics. A secondary antibody was added at the end of the experiment to clearly differentiate between non-internalized and internalized targets, illustrating the highly phagocytic nature of these cells (**Figure 4C**). While these bead-based assays allow for an accurate measurement of macrophage phagocytic activity, live cell imaging of fluorescently-labeled pathogens can be used to track their rate of uptake – and if imaged for a sufficient period of time – pathogen degradation can be also be quantified as a loss of fluorescent puncta (**Figure 4D**). Following engulfment, phagocytosed pathogens undergo a vesicular trafficking-mediated process that terminates in the fusion of lysosomes to the phagosome, thus delivering the vacuolar ATPase that acidifies the phagosome and the degradative enzymes that kill and degrade the pathogen ^27^. The fusion of lysosomes to pathogen-mimic-containing phagosomes was quantified using live-cell imaging of macrophages whose lysosomes were labeled with fluorescent dextran (**Figure 4E**), while pathogen mimics labeled with pH-sensitive and pH-insensitive fluorophores were used to measure the pH of maturing phagosomes by ratiometric imaging (**Figure 4F-G**). By multiplexing six assays into a single chamber and using three imaging modalities for data collection, this chamber design enabled the quantification of multiple stages of the phagocytic process using only 25,000 cells, thus allowing this experiment to be completed in just 4 hours. In comparison, our previous studies using similar approaches often required days of imaging to achieve a similar analysis of the phagocytic process ^28^.

### Easy Implementation of Diverse Chamber Designs

One advantage offered by FDM printing over other laboratory-adoptable manufacturing processes is the relative ease in which chambers with multiple parts, complex geometries, or interior structures can be created. Moreover, FDM printing can be used to produce moulds for more traditional chamber assemblies such as PDMS-cast chambers. To highlight these capacities, we designed moulds for both parallel-plate flow assays and chemotaxis chambers, using a simple lab-made oxygen plasma generator to attach the PDMS chambers to glass coverslips (**Figure 5A-B**) ^29^. Using the flow chamber, we replicated our previous quantification of the avidity of the apoptotic cell binding receptor MER-tyrosine kinase (MERTK) by measuring the binding of apoptotic cell mimics under increasing shear. This experiment produced results identical to those we reported previously, when performing the same experiment with a commercial flow chamber (**Figure 5C**) ^30^. The chemotaxis of human neutrophils to the bacterial peptide formyl-Methionine-Leucine-Phenylalanine was assessed using the chemotaxis chamber, producing chemotactic movement with directionality and speed similar to what we have reported previously using the classical under-agarose chemotaxis assay (**Figure 5D-F**) ^31,32^. The minimum feature size achievable in these PDMS casts, in theory, should equal the nozzle diameter and z-resolution of the printer. In practice, on our system we observe an ±8% variation in the width of print-lines, likely due to slight variations in the filament extrusion rate and/or the formation of pressure ridges between neighbouring print lines. This limits the practical minimum feature size to objects 2-3 print lines in width (**Figure 5G**). To illustrate the ability of FDM printing to produce complex chambers, we designed a reusable magnetic Leiden chamber designed to fit 18 mm circular coverslips. Using these Leiden chambers, we performed phagocytosis assays with transgene-expressing macrophages (**Figure 5H-I**), again producing results equivalent to those we have previously reported ^33^.

**Figure 5:**
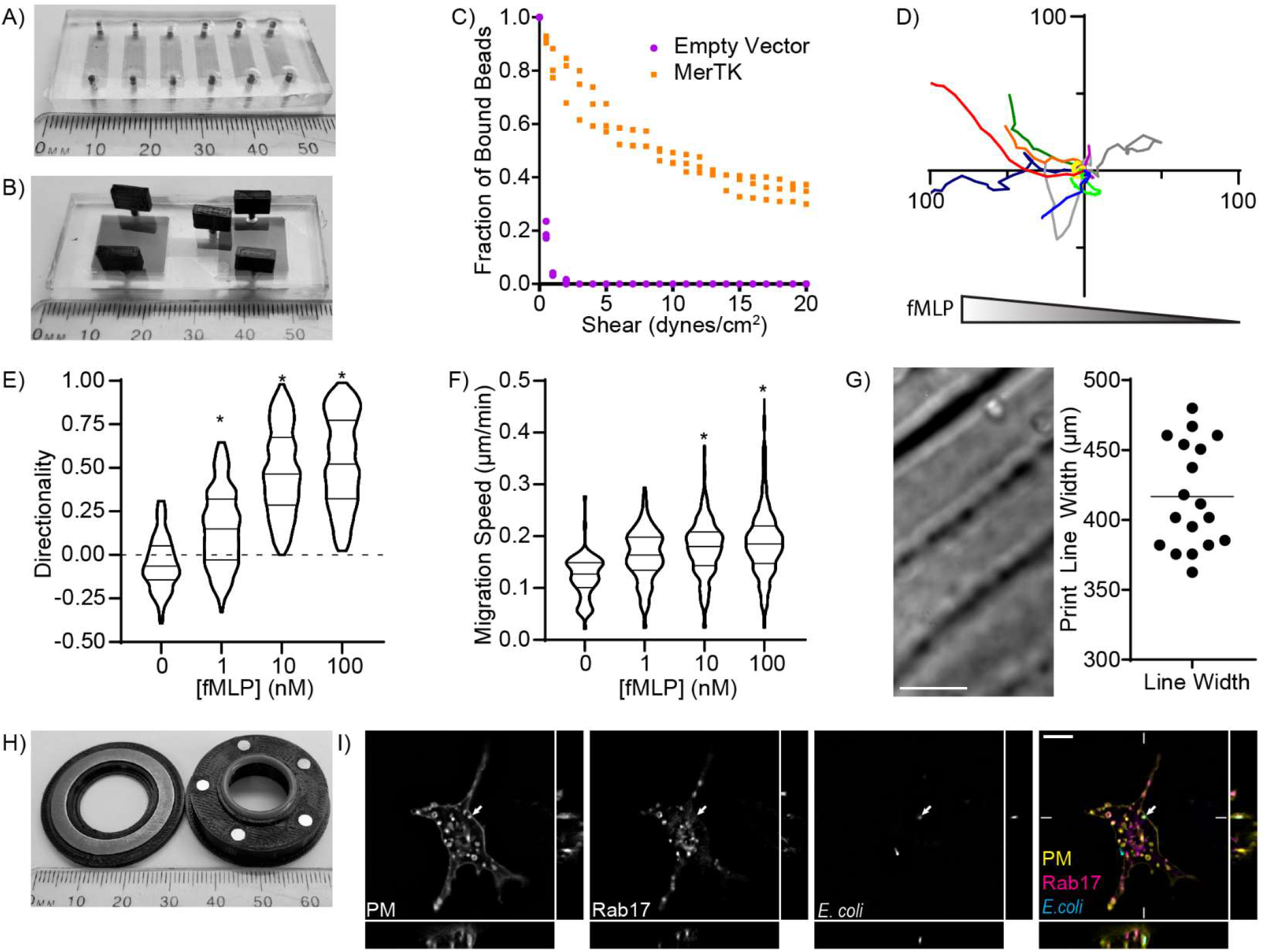
Broad Applicability of FDM Printing for Chamber Design. **A-B)** Fully assembled parallel-plate flow chamber (A) and chemotaxis chamber (B) produced as PDMS casts from FDM printed moulds. Chambers are filled with a coloured fluid to emphasize their internal structure. **C)** Adhesion of apoptotic cell mimics by MerTK-expressing and non-expressing (Empty Vector) Cos7 cells exposed to increasing shear rates in the PDMS flow chamber. Data plots 3 independent experiments. **D-F)** Chemotaxis of DRAQ5-labeled human neutrophils, in the PDMS chemotaxis chamber, showing migration tracks of 10 randomly selected cells moving towards 10 nM fMLP (D), and the directionality (E) and speed (F) of chemotaxis to increasing concentrations of fMLP. Data is representative of (D) or quantifies (E-F) a minimum of 240 migration tracks collected over 3 independent experiments. * = p < 0.01 compared to 0 nM, Kruskal-Wallace with Dunn correction. **G)** DIC image (left) and quantification (right) of the variation in width of print-lines, as moulded into a PDMS cast of a mould printed with a 0.4 mm diameter nozzle. **H)** Image of an FDM-printed magnetic Leiden chamber. **I)** Z-stack of a RAW264.7 macrophage expressing a plasma membrane marker (PM-GFP) and mCherry-Rab17 which has phagocytosed labeled *E. coli*. Phagocytosed E. coli can be found in a plasma-membrane derived vacuole (arrow) free of the efferosome marker Rab17. Positioning of the x/y/z-projections are indicated by the short lines in the colour image, scale bar is 10 μm. Image is representative of 30 cells imaged in 3 independent experiments.

## Discussion

In this study, we have demonstrated that FDM printing can be used to create imaging chambers of a variety of designs suitable for live- and fixed-cell microscopy across a range of imaging modalities. Compared to machining and photolithography approaches, FDM printing can be employed with relatively inexpensive equipment that requires minimal expertise to operate. Moreover, the per-chamber cost of this approach is low – often less than 5% the cost of an equivalent commercial chamber – with print times for the average chamber taking less than an hour. The compatibility across a range of imaging modalities, design flexibility, low cost, speed of manufacture, and ease of implementation makes the use of FDM printed imaging chambers an approach that can be adopted by many laboratories.

While the manufacture of imaging chambers by FDM printing offers many benefits, the flexibility of chamber design and compatibility with multiple imaging modalities including super-resolution microscopy, are the greatest strengths of this approach. Specifically, the ability to incorporate a #1.5 (0.17 mm thick) coverslip made of glass with a refractive index of 1.515 is a key strength, as the high-numerical aperture objective lenses required for many microscopy modalities are built assuming a coverslip of this thickness and refractive index will be placed between the lens and the subject ^2,3^. Deviating from this thickness and refractive index can profoundly affect image quality, with high-numerical aperture lenses suffering a >80% loss in light gathering power and the creation of significant spherical and chromatic aberration by coverslips deviating by as little as 0.02 mm in thickness ^2^. These issues are compounded by plastic coverslips, which, in addition to imperfect matching of thickness and refractive index, also have higher rates of light scattering, autofluorescence, and can interfere with imaging modalities requiring polarized light ^34^. In addition to providing ideal imaging conditions, the ease in which FDM chambers can be designed and prototyped, and the ability to design chambers with complex geometries, allows for chambers to be optimized to the specific needs of an experiment.

A major concern with using FDM printed imaging chambers is the potential for cell-incompatible materials to be present in the completed chamber. Surprisingly, none of the plastics we tested showed any overt signs of cell toxicity. Given our results and the success of other groups ^7,15,16,18^, we recommend using PLA for the production of imaging chambers. While we have tested a range of PLA filaments without any signs of cell toxicity, natural (e.g. unpigmented) or black pigmented filaments are preferred as many coloured filaments have high autofluorescence and are therefore not suitable for fluorescence microscopy (data not shown). A second concern with FDM printed chambers is sanitizing or sterilizing the chambers. FDM printing takes place at elevated temperatures (190° C or higher), meaning that completed prints should be free of vegetative bacteria, viruses, fungi, and most spores ^35^. However, assembling the chambers without introducing contamination can be challenging. Herein, and in the studies of others ^7,16^, a short (5-15 minute) immersion in 70% ethanol was sufficient to allow for cell culture without contamination. Indeed, we were able to culture both HeLa cells and J774.2 macrophages for five days without overt signs of infection in antibiotic-free medium using chambers sanitized in this fashion. Unfortunately, none of the plastics tested in this study survived steam autoclaving (data not shown), but complete sterilization of PLA and nylon should be possible using ethylene dioxide ^18,36^.

An important consideration when FDM printing imaging chambers is optimizing printing parameters to produce a chamber with the necessary physical characteristics. FDM prints are typically built as surfaces (perimeters) made of one or two layers of solid plastic, with the interior space filled by a 3D pattern (infill) that is largely empty space. While this approach saves on material and speeds printing, the large amount of air trapped in the infill acts as an insulator, slowing temperature equilibration when chambers are placed into an incubator or onto the heated stage of a microscope. Of greater concern, FDM printing creates chambers with porous walls that are permeable to gasses, and to a lesser extent, permeable to culture medium. This porosity is an issue with FDM produced parts that cannot be eliminated through changes to printing parameters alone (e.g. printing temperature, layer height, etc) ^37,38^. The primary printing parameter that affects porosity is the flow rate – e.g. how much plastic is extruded relative to the amount of plastic estimated to be required to fill the print volume, with flow rates of 0.98 to 1.0 – which are the default for most printers – producing the lowest porosity ^38^. While our work demonstrated that increasing wall thickness to 1.5 mm (e.g. 4 perimeters with a 0.4 mm print head) could reduce porosity to a level compatible with most experimental needs, the complete elimination of porosity requires alternative approaches such as solvent smoothing or treatment with a sealant. Unfortunately, both solvent smoothing and sealants can alter the dimensions of printed parts ^37,39^, and smoothing PLA requires the use of toxic solvents such as tetrahydrofuran ^40^. While higher printing temperatures might be expected to lower porosity by promoting better bonding between layers, experimental evidence indicates that the print temperature does not meaningfully affect porosity ^38^, while elevated printing temperatures can produce cytotoxic materials such as polyacrylic acid through the thermal decomposition of the printer filament ^41^. To our knowledge, PDMS remains the only available cell-compatible adhesive and should be used for assembly of any components that will directly contact cells or culture medium ^22^. However, cyanoacrylate (instant) glues are an excellent choice for assembling portions of chambers that will not be exposed to cells.

Minimum feature size and the precision with which small features can be formed is a limitation of FDM printing, both for prints used directly as chambers and for prints used as moulds for casting PDMS. Consistent with the observations of others, our results indicate that the spatial variation intrinsic to FDM printing allows for larger (e.g. millimetre-scale) structures to be formed accurately, but not structures near the size of the nozzle diameter or z-resolution of the printer ^42,43^. If higher precision is required, stereolithography (SLA) printing may be a superior option to FDM printing. In SLA printing, a laser or backlit LCD screen is used to induce the polymerization of a photosensitive resin, with objects built by the sequential addition of photopolymerized layers ^44^. SLA printing has much higher resolution than FDM printing (50 μm or better), and while some distortion can occur during polymerization, these deviations are much smaller than the variation we observed in our FDM prints ^45^. Unfortunately, common SLA resins are toxic, and therefore “desktop” SLA printers are currently limited to producing moulds for PDMS casting ^46^.

Herein, we have illustrated the ease with which inexpensive FDM printed imaging chambers can be designed to fulfill a broad range of imaging-based experiments. While our chambers were relatively simple, complex features allowing for experimental conditions to be directly manipulated by the chamber (e.g. stimulating electrodes and valves), as well as equipment to provide experimental readouts (e.g. temperature sensors) can be directly incorporated into these designs ^7,47^. Moreover, we have shown that FDM printing can be used to create moulds for preparing conventional PDMS microfluidic chambers, thus offering additional design options and a low-cost entry point to PDMS microfluidics. Because our approach incorporates glass coverslips into the imaging chambers, these chambers are compatible with demanding microscopy methods that require high numerical aperture optics. The combination of design flexibility and optimal optical characteristics overcomes the limitations of many existing imaging chambers, thus offering new imaging opportunities across the life sciences.

## Materials and Methods

### Materials

MerTK, Lifeact-RFP, mCherry-Rab17 and PM-GFP constructs were prepared previously ^30,33,48^. #1.5 thickness glass coverslips, polydimethylsiloxane (PDMS), 16% paraformaldehyde (PFA) and glass slides were from Electron Microscopy Sciences. J774.1 macrophages and HeLa cells were from Cedarlane Labs. DMEM, RPMI, fetal bovine serum (FBS), and trypsin-EDTA were from Wisent. M-CSF was from Peprotech. Microbeads were from Bangs Laboratories. 1-palmitoyl-2-oleoyl-sn-glycero-3-phosphatidylserine, biotinylated phosphatidylethanolamine, and 1-palmitoyl-2-oleoyl-sn-glycero-3-phosphatidylcholine were from Avanti Polar Lipids. All cell culture plastics and plastic consumables, Neon transfection system, pHrodo, NBT, TRITC-Dextran, Permafluor mounting medium, phalloidin-AlexaFluor-555, wheat germ agglutinin-AlexaFluor-647 and Hoechst 33342 were from ThermoFisher Canada. Rat IgG was from Sigma-Aldrich. Acetate tape (“Magic tape”, Scotch brand) was from Staples Office Supply. GenJet Plus transfection reagent was from FroggaBio. μ-Slide 8-well chambered polymer coverslips were from Ibidi. Prism 8 software was from GraphPad (La Jolla, California). Matlab was from Mathworks. FIJI was downloaded from https://fiji.sc/ ^49^. A Prusa i3 MK3S 3D Printer was purchased from Prusa research (Czech Republic). 1.75 mm diameter 3D printing filament (TRUE Food Safe PLA, TRUE Food Safe PETG, Nylon 645 and Polymax PC) was from Filaments.ca. All other materials were purchased from Bioshop Canada (Burlington, Canada).

### Cell Culture

COS7, J774.2 and RAW264.7 macrophages were cultured in DMEM + 10% FBS in a 5% CO_2_/37°C incubator. J774’s and RAW264.7 cells were split upon reaching 80% confluency by scraping the cells into suspension and then diluting 1:10 into fresh media. HeLa cells were cultured in RPMI+ 10% FBS in a 5% CO_2_/37°C incubator; HeLa ad COS7 cells were split upon reaching confluency by washing the cells once with phosphate buffered saline (PBS, 137 mM NaCl and 10 mM Na_2_HPO_4_), detached with Trypsin-EDTA, and diluted 1:10 in fresh media prior to replating. For experiments, cells were plated at a density of 1,000 cells/mm^2^ into 3D printed chambers, Ibidi μ-Slide 8-well chambered polymer coverslips, or onto 18 mm circular coverslips placed into the wells of a 12 well plate. For transfection-based experiments, HeLa or Cos7 cells on 18 mm coverslips were transfected with 0.75 μg of DNA and 2.5 μL of GenJet Plus DNA In Vitro Transfection Reagent as per the manufacturer’s instructions 24 hrs prior to imaging, whereas RAW264.7 cells were transfected with 5 μg DNA using a Neon electroporation system as per the manufacturer’s instructions. Transfections in FDM-printed chambers were conducted in the same manner, adjusting the quantity of DNA and transfection reagent to maintain a consistent quantity of DNA per area of transfected cells. For immunolabeling, cells were fixed for 15 min at 37°C with 4% paraformaldehyde in PEM buffer (80 mM PIPES pH 6.8, 5 mM EGTA, 2 mM MgCl_2_, ^50^), permeabilized with and blocked for 1 hr with PEM + 0.1 Triton X-100 + 1% bovine serum albumin, and labeled as indicated below.

### Microscopy and Image Analysis

All microscopy was performed using a Leica DMI6000B microscope equipped with 40×/1.30 NA, 63×/1.40NA and 100×/1.40 NA objectives, photometrics Evolve-512 delta EM-CCD camera, heated/CO_2_ perfused stage, Chroma Sedat Quad filter set with blue (Ex: 380/30, Em: 455/50), green (Ex: 490/20, Em: 525/36), red (Ex: 555/25, Em: 605/52) and far-red (Ex: 645/30, Em: 705/72) filter pairs, and the LAS-X software platform with Live Data Mode. Image analysis was performed in FIJI ^49^ or in Matlab.

### SRRF Microscopy

HeLa or J774.2 cells were fixed as above and labeled with Wheat Germ Agglutinin-AlexaFluor-647 as per the manufacturer’s instructions. The cells were then permeabilized and stained for 10 min at room temperature with 1 μg/mL Hoechst 33342 and a 1:1,000 dilution of phalloidin-AlexaFluor-555. 250 images of each channel were captured, adjusting the exposure intensity, exposure time, and EM gain to minimize the acquisition time (typically 50 ms/image). The resulting images were exported as TIFF stacks and reconstructed using the NanoJ-SRRF plugin in FIJI, using a ring radius of 0.5, radiality magnification of 5, and 6 ring axes ^51^. The resolution of the resulting SRRF images was determined using a line profile of single actin fibrils in FIJI. The resulting intensity values were imported into Prism, the curve matching tool used to match the line profile to a Gaussian curve, and the full width at half maximum calculated. A minimum of three line profiles were collected per image, from a minimum of 10 images per experiment.

### Cell Toxicity Assays

Crystal violet was used to quantify cell viability using the method of Feoktistova *et al* ^52^. FDM printed plastic rings (outer ⌀: 21.4 mm, inner ⌀: 19.4 mm) were made in various plastics using the manufacturer’s recommended printing speeds, temperature and fan settings. The final prints were cleaned and sanitized by soaking in 70% ethanol for 1 hr, dried in a biosafety cabinet, and then briefly soaked in sterile PBS to remove residual ethanol. Rings were placed into the wells of a 12-well plate and 5 × 10^4^ cells in 1 mL of media added to each well. Cells were grown for 5 days with no media exchange. The rings were then removed, the wells washed with PBS and stained with 0.5 mL of a 0.5% w/v crystal violet, 25% v/v MeOH solution on an orbital shaker for 30 min at room temperature. The plate was washed by repeated immersion in water, using fresh water for each immersion, until no colour was visible in the wash. The plate was dried for 1 hr at room temperature, and then decolorized by adding 1 mL of a 2% w/v SDS, 50% v/v MeOH solution for 20 min with vigorous shaking at room temperature. Crystal violet incorporation into viable cells was quantified by measuring the OD595 with an Eon microplate reader. Absorbance values were calculated as a mean of 25 reads per well, spaced out equally in a 5 × 5 grid.

To identify non-cytotoxic effects, cell morphology was quantified for cells cultured for 48 hrs on 18 mm alone or with the 3D printed rings described above. As a positive control, cells were treated with 1 μM staurosporine to induce apoptosis. Next, cells were fixed for 20 min in 4% PFA and stained with AlexaFluor555-labeled phalloidin, AlexaFluor-647 labeled wheat germ agglutinin, and counterstained with Hoechst, as described above. The samples were then mounted on slides with Permafluor and imaged with SRRF microscopy. The resulting images were imported into Matlab and the morphology of the cells quantified using adjacency statistics, performing adjacency statistics using thresholds of the mean and the mean ± 1.5 standard deviations, thus producing a 54-dimenional “fingerprint” of the cells morphology ^19^. Principal component analysis was then used to reduce the 54-dimensional dataset to two dimensions. Cytotoxicity of cyanoacrylate glue was assessed using RAW264.7 cells expressing a plasma membrane marker that were placed into a chamber whose coverslip was attached by cyanoacrylate glue or PDMS. Cell death was identified via blebbing of the plasma membrane during time-lapse microscopy.

### 3D Printing

Chambers were designed using TinkerCAD (www.tinkercad.com), or the free academic versions of Autodesk Netfab and AutoCAD. Chamber designs were exported in the stereolithography (.STL) file format and sliced onto 3D printer compatible G code using Ultimaker CURA 4.0 software. Unless otherwise noted, all chambers were sliced using 0.1 mm layer height, 100% infill, using the following printing temperature, bed temperature and fan settings, respectively: PLA – 210°C/60°C/100% fan, PETG – 240°C/85°C/40% fan, PC: 270°C/115°C/0% fan, nylon – 245°C/110°C,100% fan. All prints were made on a Prusa i3 MK3S 3D Printer equipped with a 0.4 mm nozzle, with a thin layer of polyvinyl acetate added to the print bed to improve print adhesion. Prints were cleaned with isopropyl alcohol. STL files for all chambers used in this assay are available in the supplemental materials.

### Chamber Assembly

3D prints were attached to coverslips using PDMS. 900 μl of PDMS was placed into a 1.7 mL microcentrifuge tube, and then mixed with 100 μL of the curing agent. After mixing, the solution was degassed by a 30 sec, 21,000 × g centrifugation. A thin channel, roughly 1.5 times the width and length of the print was made on a glass plate using a double-thickness of acetate tape for the walls. The PDMS was then spread in this channel, creating a layer of unpolymerized PDMS ~0.15 mm thick. The print was pressed into the PDMS, and then placed onto a coverslip. The assembled chamber was then placed on a hot plate set to 60°C for 2 hrs, then at room temperature overnight, to fully polymerize the PDMS. For assembly with cyanoacrylate glue, small droplets of glue were placed on the FDM printed part, spaced equally around the coverslip-print interface. A coverslip was then placed onto the glued surface, and the glue allowed to cure for 24 hr before use. All chambers were sanitized by soaking them in 70% ethanol for 1 hr and then dried in a biosafety hood, followed by a final soak in sterile PBS.

### Pressure and Vacuum Testing

A pressurizable test chamber was printed with 0.5 cm thick walls, a 0.5 cm^3^ interior volume, with a 10 mm × 10 mm opening. An 18 mm, #1.5 thickness square coverslip was sealed over this opening with PDMS and the chamber connected via a Luer adaptor to a custom-build pressure/vacuum apparatus with a Honeywell high-sensitivity pressure sensor with real-time data logging. For pressure testing, the test chamber and connecting line were filled with water and pressurized using nitrogen. Chamber pressure was increased in 50 kPa increments, with the chamber monitored for leakage for 30 sec between increments. Chambers were pressurized until they failed. For vacuum testing, the test chamber and connecting lines were flushed with nitrogen, and then a vacuum was applied using a ThermoFisher LAV3G high vacuum pump. Chambers were evacuated to ~1 kPa (~99.9% vacuum), and the pressure monitored for 5 min.

### Flow Chamber Analyses

To determine the accuracy of FDM chamber construction, a flow chamber was designed bearing dual flow areas 16 mm × 2.5 mm × 1.5 mm (L×W×H) in size and printed in PLA using layer heights of 0.07 mm to 0.2 mm. Prior to assembly, the width and depth of the chambers were measured with a high-resolution (0.02 mm accuracy) digital caliper. Time lapse images were captured at 40× magnification and 10 frames-per-second, as 5 μm diameter silica beads, suspended as a 1:10,000 dilution in distilled water, were perfused through the chamber at 0.054 and 0.108 mL/s using a 3.0 mL syringe and a syringe pump in draw mode. The microscope was focused as deeply into the chamber as possible to limit the effects of wall shear on bead velocity. FIJI was used to calculate the displacement of beads between time points, from which bead velocity was determined. Only beads present in the field of view for at least 3 successive frames were used to calculate velocity, measuring the average velocity across time points. The fluid velocity of a chamber was defined as the average velocity of a minimum of 50 beads.

### 12CA5 Fab generation and labeling

Media was collected from 12CA5 hybridoma cells cultured for 5 days at high density (~5×10^8^/mL) in serum-free hybridoma media. The media was cleared with a 1,500×g/30 min centrifugation and then concentrated with a 60 kDa cut off centrifuge concentrator. The concentrate was diluted to 5 mg/ml in PBS plus 10 mM EDTA and 20 mM cysteine-HCl, and a 50% volume of immobilized papain added. After an 18 h/37°C incubation the papain was then removed by a 1000×g, 15 min centrifugation and Fab fragments separated using fast protein liquid chromatography on a Sephacryl S100 column. Aliquots corresponding to Fab fragments were pooled and labeled with a Cy3 labeling kit (Abcam) as per the manufacturer’s instructions. Labeled Fab fragments were diluted to 1 mg/ml and stored frozen in PBS + 20% glycerol.

### Single Molecule Localization Microscopy

Single molecule localization microscopy was performed as described previously ^53,54^. Briefly, HeLa cells were split into 3D printed imaging chambers or Ibidi μ-Slide 8 plastic-bottomed chambers and transfected with HA-tagged CD93 using GenJet as per the manufacturer’s instructions. 24 hrs later, the cells were cooled to 10°C and the CD93 labeled by incubating with a 1:10,000 dilution of Cy3-labeled 12CA5 Fab fragments in PBS for 10 min. The cells were then washed 3× with PBS, placed in imaging buffer (150 mM NaCl, 5 mM KCl, 1 mM MgCl_2_, 100 mM EGTA, 2 mM CaCl_2_, 25 mM HEPES, and 1500 g/L NaHCO_3_, pH 7.4) and transferred to the heated/CO_2_ perfused stage of our microscope. Time-lapse acquisitions of the basolateral side of the cells were captured, imaging at 10 frames/s for 30 s. The resulting videos were cropped to the area containing the cell, and the image sequences imported into Matlab. Individual fluorophores were identified and resolved at super-resolution using the mixed model Gaussian fitting algorithm of Jaqaman *et al* ^24^, and both the estimated full-width at half-maximum of the images point-spread function (in pixels), and the precision of fluorophore localization, exported for subsequent analysis.

### Phagocytosis Assays

Phagocytosis was quantified as described previously ^33^. Briefly, J774.2 macrophages were split into the multiplex imaging chamber or onto 18 mm circular coverslips and allowed to recover for 24 hrs. Where required, the cells were transfected with mCherry-Rab17 and PM-GFP using Lipofectamine 2000 as per the manufacturer’s instructions. Phagocytic targets were generated by 1) incubating 10 μL of 5 μm diameter silica beads with 100 μL of unlabeled 1μg/mL rat IgG, 2) incubating 10 μL of 5 μm diameter silica beads with 100 μL of unlabeled 1μg/mL rat IgG plus 1:2,000 dilutions of AlexaFluor-647 labeled rat IgG ± 1:2,000 dilution of pHrodo labeled rat IgG, or 3) incubating 1 × 10^6^ E. coli K12 in PBS with a 1:2,000 dilution of Cell Proliferation Dye eFluor 670, for 30 min at room temperature. Where required, macrophages were pre-loaded with 100 μg/ml TRITC-conjugated dextran for 16 hrs, followed by a 90 min chase with serum-free DMEM. For live-cell imaging, the multiplex imaging chamber was transferred to the heated/CO_2_ perfused stage of our microscope, and individual acquisition parameters set for each well using Live Data Mode. Phagocytic targets were added at a 10:1 (target: macrophage) ratio and the samples imaged every 1 min for 2 hrs using the 60× objective. For end-point assays, cells were washed once with PEM buffer and then fixed for 15 min with 4% PFA in PEM, and non-internalized phagocytic targets labeled using a Cy3-labeled goat-anti-rat Fab fragment, and the sample imaged using the 60× objective. Spontaneous migration was quantified using the Manual Tracking plugin in FIJI ^49^, phagosome-lysosome fusion was quantified by identifying co-localization between TRITC-dextran and AlexaFluor-647 labeled phagocytic targets, and pH was quantified via the pHrodo:AlexaFluor-647 ratio and converted to pH using a standard curve generated by imaging the same bead preparation in saline buffered with 50 mM MES (pH 4.0, 5.0) or 50 mM Tris (pH 6.0, 7.0 and 7.4).

### PDMS Chamber Construction

Moulds to cast PDMS chambers were FDM printed using PLA with 100% infill, 0.1 mm layer height, with the ironing (surface smoothing) feature enabled. For each chamber, 5 mL of PDMS polymer and 500 μL of PDMS catalyst were mixed in a 14 mL snap-cap tube and degassed by centrifugation at 2,000 × g for 1 min. The mixture was poured into FDM printed chemotaxis moulds and subjected to 30 min of vacuum at ~60 kPa. The moulds were then transferred to a 37°C incubator for 24 hr. The chemotaxis chambers were freed from the mould with a scalpel, washed 3× with 70% ethanol, rinsed 3× with distilled water, and dried using a Kimwipe. The chamber and clean 25 mm × 50 mm #1.5 thickness coverslip were placed in a custom-made glass vacuum chamber which was flushed 2 times with oxygen, and then subjected to an ~1 kPa vacuum. The chamber was then exposed to 2 sec of plasma by placing the chamber in a 1250 W consumer microwave at maximum power ^29^. The chamber was then pressed against the coverslip, and the assembled chamber placed on a hot plate at 90°C for 1 hr to covalently bond the chamber to the coverslip.

### Apoptotic Mimic Capture Assay

PDMS flow chambers consisting of 6 parallel channels 5 mm × 15 mm × 0.2 mm with 1 mm dimeter connection ports were cast and attached to 25 mm × 50 mm coverslips as described above. Cos7 cells were transfected with empty vector or a vector expressing human MerTK-GFP. 24 hr later, the Cos7 cells were detached using trypsin and seeded at 75% confluency into assembled flow chambers and allowed to recover in a 37°C/5% CO_2_ incubator for an additional 18-24 hr. Apoptotic cell mimics were generated by adding 10 μL of 3 μm diameter silica beads to a mixture of 3.2 μmol phosphocholine and 0.8 μmol phosphatidylserine in chloroform, and dried under nitrogen gas. The dried beads were suspended in 1 mL of PBS and washed three times by centrifuging for 1 min at 4,500 × g and resuspending in 100 μL of PBS. After washing, 3 μL of beads were suspended in 1 mL of RPMI + FBS, perfused into the flow chamber, and incubated for 10 min. The chamber was then mounted on the heated stage of our microscope and attached to a syringe pump in draw mode via silicone tubing connected to 16-gauge blunted needle. The same tubing/needle setup was used to connect the other side of the flow chamber to a reservoir of media 37°C media. The sample was imaged using DIC to identify bound beads and by fluorescence microscopy (Ex: 490, Em: 525) to identify transfected cells. The sample was then subjected to 30 sec of shear at 0.5 dynes/cm^2^, and the DIC imaging was repeated. This shear and DIC imaging process were repeated for 1-20 dynes/cm^2^, in 1 dynes/cm^2^ increments. MERTK-expressing cells were then identified in the fluorescence image, and the fraction of initially bound beads were quantified for each cell at each shear rate. A minimum of 30 transfected cells were imaged for each condition.

### Chemotaxis Assay

PDMS chemotaxis chambers consisting of two 600 μL reservoirs connected by a 5 mm × 10 mm × 0.2 mm channel were cast and attached to 25 mm × 50 mm coverslips as described above. The collection of blood from healthy donors was approved by the Health Science Research Ethics Board of the University of Western Ontario and venipuncture was performed in accordance with the guidelines of the Tri-Council Policy Statement on human research. 8 mL of blood was drawn into a heparinized vacuum tube and layered over an equal volume of Lympholyte Poly, followed by a 300 × g/35 min centrifugation. The neutrophil band was removed, diluted to 50 mL with PBS and pelleted using 300 × g/5 min centrifugation. The neutrophils were suspended in 1 mL of PBS, stained for 5 min with 1:2,000 DRAQ5, then washed twice with 1 mL of PBS and 300 × g/5 min centrifugations and suspended in RPMI + 10 % FBS at 2 × 10^6^/mL. One reservoir was used as a “sink” and filled with 600 μL of RPMI + 10 % FBS, and the loading ports closed with FDM printed plugs. 50 μL of the neutrophil suspension was added to the channel, the loading port closed, and the second reservoir filled with 600 μL of fMLP in RPMI + 10% FBS. The chamber was placed on the heated stage of our microscope and imaged at 40× magnification using fluorescence microscopy (Ex: 645, Em: 705), with images captured every 30 sec for 4 hrs. The resulting time-lapse sequences were then analysed for migration speed and directionality using the TrackMate plugin in FIJI ^49,55^.

### Statistics

Unless otherwise noted, data are presented as mean ± SEM. All statistical analyses were performed in Graphpad Prism, using α = 0.05 as a significance cut-off.

## Acknowledgements

This study was funded by a Grant-in-Aid from the Ontario Lung Association and an Early Researcher Award from the Ontario Ministry of Research, Innovation and Science to BH. The funding agencies had no role in study design, data collection and analysis, decision to publish, or preparation of the manuscript.

## Conflict of Interest

The authors declare no financial or commercial conflict of interest.

## Notes

#### Summary of Updates

The manuscript has been re-focused to concentrate more on the multiplex and multimodal microscopy opportunities offered by this method. Additional data highlighting these capabilities is provided, as is additional descriptions of the strengths and weaknesses of the approach. Additional details on some of the analyses and reagents we used have also been included.

## References

1. Lee, J.-Y. & Kitaoka, M. A beginner’s guide to rigor and reproducibility in fluorescence imaging experiments. MBoC 29, 1519–1525 (2018).

2. Inoue, S. & Oldenbourg, R. Microscopes, Chap. 17 in Handbook of Optics, Vol. II, M. Bass, Ed. (McGraw-Hill, New York, 1995).

3. Abramowitz, M., Spring, K. R., Keller, H. E. & Davidson, M. W. Basic principles of microscope objectives. BioTechniques 33, 772–774, 776–778, 780–781 (2002).

4. Phillipson, M. et al. Intraluminal crawling of neutrophils to emigration sites: a molecularly distinct process from adhesion in the recruitment cascade. J. Exp. Med. 203, 2569–2575 (2006).

5. Khan, A. I., Heit, B., Andonegui, G., Colarusso, P. & Kubes, P. Lipopolysaccharide: a p38 MAPK-dependent disrupter of neutrophil chemotaxis. Microcirculation 12, 421–432 (2005).

6. Turek, M., Besseling, J. & Bringmann, H. Agarose Microchambers for Long-term Calcium Imaging of Caenorhabditis elegans. JoVE (Journal of Visualized Experiments) e52742 (2015) doi:10.3791/52742.

7. Wardyn, J. D., Sanderson, C., Swan, L. E. & Stagi, M. Low cost production of 3D-printed devices and electrostimulation chambers for the culture of primary neurons. J. Neurosci. Methods 251, 17–23 (2015).

8. Alessandri, K. et al. All-in-one 3D printed microscopy chamber for multidimensional imaging, the UniverSlide. Sci Rep 7, 42378 (2017).

9. Wang, Y.-X. et al. A multi-component parallel-plate flow chamber system for studying the effect of exercise-induced wall shear stress on endothelial cells. BioMedical Engineering OnLine 15, 154 (2016).

10. Sollier, E., Murray, C., Maoddi, P. & Di Carlo, D. Rapid prototyping polymers for microfluidic devices and high pressure injections. Lab Chip 11, 3752–3765 (2011).

11. Hwang, Y. & Candler, R. N. Non-planar PDMS microfluidic channels and actuators: a review. Lab Chip 17, 3948–3959 (2017).

12. Leclerc, E., Sakai, Y. & Fujii, T. A multi-layer PDMS microfluidic device for tissue engineering applications. in IEEE The Sixteenth Annual International Conference on Micro Electro Mechanical Systems, 2003. MEMS-03 Kyoto 415–418 (2003). doi:10.1109/MEMSYS.2003.1189774.

13. Henley, W. H., Dennis, P. J. & Ramsey, J. M. Fabrication of microfluidic devices containing patterned microwell arrays. Anal. Chem. 84, 1776–1780 (2012).

14. Salentijn, G. I., Oomen, P. E., Grajewski, M. & Verpoorte, E. Fused Deposition Modeling 3D Printing for (Bio)analytical Device Fabrication: Procedures, Materials, and Applications. Anal. Chem. 89, 7053–7061 (2017).

15. Rimington, R. P., Capel, A. J., Christie, S. D. R. & Lewis, M. P. Biocompatible 3D printed polymers via fused deposition modelling direct C2C12 cellular phenotype in vitro. Lab Chip 17, 2982–2993 (2017).

16. Gulyas, M., Csiszer, M., Mehes, E. & Czirok, A. Software tools for cell culture-related 3D printed structures. PLoS ONE 13, e0203203 (2018).

17. Tyson, A. L., Hilton, S. T. & Andreae, L. C. Rapid, simple and inexpensive production of custom 3D printed equipment for large-volume fluorescence microscopy. Int J Pharm 494, 651–656 (2015).

18. Athanasiou, K. A., Niederauer, G. G. & Agrawal, C. M. Sterilization, toxicity, biocompatibility and clinical applications of polylactic acid/polyglycolic acid copolymers. Biomaterials 17, 93–102 (1996).

19. Hamilton, N. A., Pantelic, R. S., Hanson, K. & Teasdale, R. D. Fast automated cell phenotype image classification. BMC Bioinformatics 8, 110 (2007).

20. Tymrak, B. M., Kreiger, M. & Pearce, J. M. Mechanical properties of components fabricated with open-source 3-D printers under realistic environmental conditions. Materials & Design 58, 242–246 (2014).

21. Ouellet, E., Yang, C. W. T., Lin, T., Yang, L. L. & Lagally, E. T. Novel carboxyl-amine bonding methods for poly(dimethylsiloxane)-based devices. Langmuir 26, 11609–11614 (2010).

22. Bélanger, M. C. & Marois, Y. Hemocompatibility, biocompatibility, inflammatory and in vivo studies of primary reference materials low-density polyethylene and polydimethylsiloxane: a review. J. Biomed. Mater. Res. 58, 467–477 (2001).

23. Oda, T., Namba, K. & Maéda, Y. Position and orientation of phalloidin in F-actin determined by X-ray fiber diffraction analysis. Biophys. J. 88, 2727–2736 (2005).

24. Jaqaman, K. et al. Robust single-particle tracking in live-cell time-lapse sequences. Nat. Methods 5, 695–702 (2008).

25. McArdle, S. et al. Migratory and Dancing Macrophage Subsets in Atherosclerotic Lesions. Circ. Res. 125, 1038–1051 (2019).

26. Flannagan, R. S., Jaumouillé, V. & Grinstein, S. The cell biology of phagocytosis. Annu Rev Pathol 7, 61–98 (2012).

27. Canton, J., Khezri, R., Glogauer, M. & Grinstein, S. Contrasting phagosome pH regulation and maturation in human M1 and M2 macrophages. Mol. Biol. Cell 25, 3330–3341 (2014).

28. Flannagan, R. S., Heit, B. & Heinrichs, D. E. Intracellular replication of Staphylococcus aureus in mature phagolysosomes in macrophages precedes host cell death, and bacterial escape and dissemination. Cell. Microbiol. 18, 514–535 (2016).

29. Hui, A. Y. N., Wang, G., Lin, B. & Chan, W.-T. Microwave plasma treatment of polymer surface for irreversible sealing of microfluidic devices. Lab Chip 5, 1173–1177 (2005).

30. Evans, A. L. et al. Antagonistic Coevolution of MER Tyrosine Kinase Expression and Function. Mol. Biol. Evol. 34, 1613–1628 (2017).

31. Heit, B., Tavener, S., Raharjo, E. & Kubes, P. An intracellular signaling hierarchy determines direction of migration in opposing chemotactic gradients. J. Cell Biol. 159, 91–102 (2002).

32. Heit, B. et al. PTEN functions to ‘prioritize’ chemotactic cues and prevent ‘distraction’ in migrating neutrophils. Nat. Immunol. 9, 743–752 (2008).

33. Yin, C., Kim, Y., Argintaru, D. & Heit, B. Rab17 mediates differential antigen sorting following efferocytosis and phagocytosis. Cell Death Dis 7, e2529 (2016).

34. El-Meanawy, A., Mueller, C. & Iczkowski, K. A. Improving sensitivity of amyloid detection by Congo red stain by using polarizing microscope and avoiding pitfalls. Diagn Pathol 14, 57 (2019).

35. Darmady, E. M., Hughes, K. E. A., Jones, J. D., Prince, D. & Tuke, W. Sterilization by dry heat. J Clin Pathol 14, 38–44 (1961).

36. Gatineau, M., El-Warrak, A. O., Bolliger, C., Mourez, M. & Berthiaume, F. Effects of sterilization with hydrogen peroxide gas plasma, ethylene oxide, and steam on bioadhesive properties of nylon and polyethylene lines used for stabilization of canine stifle joints. Am. J. Vet. Res. 73, 1665–1669 (2012).

37. Mireles, J. et al. Analysis of Sealing Methods for FDM-fabricated Parts. Proceeding from Solid Free-form Fabrication Symposium 185–196 (2011).

38. Gordeev, E. G., Galushko, A. S. & Ananikov, V. P. Improvement of quality of 3D printed objects by elimination of microscopic structural defects in fused deposition modeling. PLoS One 13, (2018).

39. McCullough, E. J. & Yadavalli, V. K. Surface modification of fused deposition modeling ABS to enable rapid prototyping of biomedical microdevices. Journal of Materials Processing Technology 213, 947–954 (2013).

40. Valerga, A. P., Batista, M., Fernandez-Vidal, S. R. & Gamez, A. J. Impact of Chemical Post-Processing in Fused Deposition Modelling (FDM) on Polylactic Acid (PLA) Surface Quality and Structure. Polymers 11, 566 (2019).

41. Kopinke, F.-D., Remmler, M., Mackenzie, K., Möder, M. & Wachsen, O. Thermal decomposition of biodegradable polyesters—II. Poly(lactic acid). Polymer Degradation and Stability 53, 329–342 (1996).

42. Armillotta, A. Assessment of surface quality on textured FDM prototypes. Rapid Prototyping Journal 12, 35–41 (2006).

43. Dey, A. & Yodo, N. A Systematic Survey of FDM Process Parameter Optimization and Their Influence on Part Characteristics. Journal of Manufacturing and Materials Processing 3, 64 (2019).

44. Rebaioli, L. & Fassi, I. Preliminary Study on the Manufacturing Feasibility of Microfeatures for Microfluidics by DLP Stereolithography. in (American Society of Mechanical Engineers Digital Collection, 2019). doi:10.1115/DETC2019-97864.

45. Fuh, J. Y. H., Lu, L., Tan, C. C., Shen, Z. X. & Chew, S. Curing characteristics of acrylic photopolymer used in stereolithography process. Rapid Prototyping Journal 5, 27–34 (1999).

46. Oskui, S. M. et al. Assessing and Reducing the Toxicity of 3D-Printed Parts. Environ. Sci. Technol. Lett. 3, 1–6 (2016).

47. Gong, H., Woolley, A. T. & Nordin, G. P. High density 3D printed microfluidic valves, pumps, and multiplexers. Lab Chip 16, 2450–2458 (2016).

48. Heit, B. et al. Multimolecular signaling complexes enable Syk-mediated signaling of CD36 internalization. Dev. Cell 24, 372–383 (2013).

49. Schindelin, J. et al. Fiji: an open-source platform for biological-image analysis. Nat. Methods 9, 676–682 (2012).

50. Pereira, P. M. et al. Fix Your Membrane Receptor Imaging: Actin Cytoskeleton and CD4 Membrane Organization Disruption by Chemical Fixation. Front Immunol 10, 675 (2019).

51. Culley, S., Tosheva, K. L., Matos Pereira, P. & Henriques, R. SRRF: Universal live-cell super-resolution microscopy. Int. J. Biochem. Cell Biol. 101, 74–79 (2018).

52. Feoktistova, M., Geserick, P. & Leverkus, M. Crystal Violet Assay for Determining Viability of Cultured Cells. Cold Spring Harb Protoc 2016, pdb.prot087379 (2016).

53. Goiko, M., de Bruyn, J. R. & Heit, B. Short-Lived Cages Restrict Protein Diffusion in the Plasma Membrane. Sci Rep 6, 34987 (2016).

54. Goiko, M., de Bruyn, J. R. & Heit, B. Membrane Diffusion Occurs by Continuous-Time Random Walk Sustained by Vesicular Trafficking. Biophys. J. 114, 2887–2899 (2018).

55. Tinevez, J.-Y. et al. TrackMate: An open and extensible platform for single-particle tracking. Methods 115, 80–90 (2017).

