## Supplementary material for "Customizable Live-Cell Imaging Chambers for Multimodal and Multiplex Fluorescence Microscopy": Readme.txt

==================================
SPACESHIPS - Live Cell Imaging Chambers
==================================
INTRODUCTION:
These are STL files for 3D printing "spaceships: (Leiden or imaging chambers) used for live and fixed cell imaging on inverted microscopes. They fit an 18 mm diameter coverslip, making it simple to culture cells directly on coverslips placed in the wells of a 12-well plate. Cells can the be fixed and stained, or transferred live to the chamber, and imaged. The chamber holds ~0.75 ml of media, which is sufficient for imaging cells for over 24 hours. For long term imaging we recommend sealing a second coverslip on the top of the chamber with VALAP (1:1:1 mix of vaseline, lanolin and paraffin) to limit dehydration. These chambers can be easily modified using CAD software to add profusion ports, electrodes, or other features.

Design fits an 18 mm round coverslip, and two base-plate sizes have been included (to fit microscope stages sized for 35 mm and 42 mm dishes). A universal top is used with both sizes of base plates. Five magnets are required per chamber and are embedded into the top of the chamber, which also houses an O-ring. A steel shim is glued into the base, allowing the chamber to be assembled by placing a coverslip in the base, then the magnetic top attached.

You will also need an O-ring for each chamber; this is what forms the water-tight seal. The O-ring diameter is critical, but the thickness can vary slightly. This chamber has been designed to seal well with #1.5 thickness (0.17 mm) coverslips, but any thickness coverslip from #1 through #3 should work.
============================
FILES:
-35mm_base.stl - chamber sized to match a 35 mm diameter culture dish
-42mm_base.stl - chamber sized to match a 42 mm diameter culture dish
-Leiden_top.stl - universal top for the chambers
-MagAlignTools.stl - printable tools to help with magnet alignment
-Readme.txt - this file

============================
REQUIRED PARTS:
Cylindrical Magnets:
-5 per chamber top, plus 1 for the alignment tool. We recommend McMaster-Carr item #5862K138 (Neodymium Magnet, Magnetized Through Thickness, 1/16" Thick, 1/8" OD)

Shims:
-1 per base. We recommend McMaster-Carr item #98055A245 (Spring Steel Ring Shim, 0.5mm Thick, 22mm ID)

Silicone O-Ring:
-1 per chamber, 18 mm (0.699") outer diameter, 1.8-2.5 mm cross-sectional diameter. We recommend McMaster-Carr item #5233T58 (High-Temperature Silicone O-Ring, 2 mm Wide, 14 mm ID)

Glue: Thin instant (cyanoacrylate) glue. We recommend Loctite 401 adhesive.

============================
ASSEMBLY:
1) Print the magnet insertion tools (file prints 2 tools, but only 1 is required). Place a small droplet of glue into the depression and insert a magnet. Let the glue cure an hour before assembling any chambers. One of these tools can be used to assemble as many chambers as you wish.

2) Print the base and top chambers. We recommend black food-grade PLA as this plastic has been previously shown to be cell-safe. We recommend printing with a 0.4 mm nozzle, 0.2 mm layer thickness, 25% gyroid infill, and to print walls at least 5 perimeters thick. Smoothing the final layer is unnecessary.

3) Pickup magnets with the rod-shaped assembly tool and insert them into the corresponding insets in the base plate. Once you are certain all the magnets are aligned properly, glue them in place with cyanoacrylate glue.

4) Use a micropipette to place a small droplet (10-15 ul) of glue into one of the depressions in the magnetic. Quickly pickup a magnet with the assembly tool and press it into the glued depression. Immediately remove the tool and use a kimwipe to clean up any residual glue. Repeat for the other four magnets.

5) Clean a shim with 70% ethanol and dry with a kimwpie. Place several small droplets of glue into the ring-shaped groove in the base plate. Press the cleaned shim into the groove, then use a kimwipe to clean up any residual glue.

5) Once all glue has cured (12 to 24 hours), insert an O-ring into the groove in the upper chamber, place an 18 mm coverslip into the base plate, and assemble. The two sections should stick to each other and form a water-tight seal.

6) Chambers can be sanitized with an 70% ethanol wash followed by UV treatment. Avoid long soaks in ethanol if printing with PLA as this can distort the chamber. Chambers are reusable but should be replaced if they start to show signs of distortion, cracking, layer separation, or if cells start to behave unusually.
============================
WARRANTY:
These chambers are provided "as is", without warranty of any kind, express or implied, including but not limited to the warranties of merchantability, fitness for a particular purpose and noninfringement. You use these chambers at your own risk.
============================
ACKNOWLEDGEMENT:
These chambers were designed by Adam Tepperman and Dr. Bryan Heit of the phagocyte biology lab (www.phagocytes.ca), and were based on the original design by Ben Goodnow at UNSW (https://github.com/PRNicovich/BenlideChambers/).
