## Supplementary material for "Customizable Live-Cell Imaging Chambers for Multimodal and Multiplex Fluorescence Microscopy": Supplemental Materials.pdf

**Supplemental Materials for: Customizable Live-Cell Imaging Chambers for Fluorescence and Super-Resolution Microscopy.**

Adam Tepperman, David Jiao Zheng, Maria Abou Taka, Angela Vrieze and Bryan Heit

This compressed file contains stereolithography (STL) files for the various 3D printed chambers and PDMS chamber moulds used in this study. This document describes each chamber and provides recommended print settings for each chamber. These chambers are provided "as is", without warranty of any kind, express or implied, including but not limited to the warranties of merchantability, fitness for a particular purpose and noninfringement. You use these chambers at your own risk.

Root Folder:
This folder contains chambers which are directly used as cell imaging chambers after attachment to a coverslip.

***2x2_chamber.stl:*** A chamber which divides an 18 mm x 18 mm coverslip into 4 separate chambers. Coverslip should be attached using PDMS. Final print size is 55 mm x 25 mm x 8.5 mm. Recommended print settings:

- Material: Food-grade black PLA
- Print Temperature: Manufacture's recommended settings (typically 215C nozzle, 60C bed)
- 0.4 mm nozzle
- 0.2 mm layer thickness
- 5 perimeters
- 25% gyroid infill

***3x3_chamber.stl:*** A chamber which divides an 18 mm x 18 mm coverslip into 9 separate chambers. Coverslip should be attached using PDMS. Final print size is 55 mm x 25 mm x 8.5 mm. Recommended print settings:

- Material: Food-grade black PLA
- Print Temperature: Manufacture's recommended settings (typically 215C nozzle, 60C bed)
- 0.4 mm nozzle
- 0.2 mm layer thickness
- 5 perimeters
- 25% gyroid infill

***Luer_flow_chamber.stl:*** Prints a flow chamber with incorporated luer connectors. This chamber was used for the printing accuracy test and requires glass coverslips be attached with PDMS to both sides of the flow region. Recommended print settings:

- Material: Food-grade black PLA
- Print Temperature: Manufacture's recommended settings (typically 215C nozzle, 60C bed)
- 0.4 mm nozzle
- 0.1 mm or 0.05 mm layer thickness
- 100% infill

PDMS Folder:
This chamber contains moulds for casting PDMS microfluidic chambers. Once cast, these chambers need to be attached to coverslips using oxygen plasma bonding.

***Chemotaxis_mold.stl:*** Mould to case a PDMS chemotaxis chamber. Chamber consists of two large reservoirs connected by a thin migration channel. A 50 mm x 25 mm coverslip should be attached using oxygen plasma bonding. Cast chamber is 50 mm x 25 mm x 5 mm. Five PDMS chamber plugs are required for this chamber. Recommended print settings:

- Material: Any PLA or PETG
- Print Temperature: Manufacture's recommended settings
- 0.4 mm nozzle
- 0.1 mm or 0.05 mm layer thickness
- Ironing activated
- 100% infill

***Flow_chamber.stl:*** Mould to cast a PDMS flow chamber. Chamber consists of six parallel and independent flow areas with ~1.5 mm diameter connecting ports at either end. A 50 mm x 25 mm coverslip should be attached using oxygen plasma bonding. Cast chamber is 50 mm x 25 mm x 5 mm. This chamber can be connected to a pump using blunted needles or used as a static chamber using ports sealed with PDMS chamber plugs. Recommended print settings:

- Material: Any PLA or PETG
- Print Temperature: Manufacture's recommended settings
- 0.4 mm nozzle
- 0.1 mm or 0.05 mm layer thickness
- Ironing activated
- 100% infill

***Micro_cell_chamber.stl:*** Mould to cast a 5 x 4 (e.g. 20-chamber) array of microwells, 5 mm x 3 mm x 0.2 mm in size. Ideal for performing a large array of experiments on minute numbers of cells. Microwells are filled via ~1.5 mm diameter connecting ports at either end. A 50 mm x 25 mm coverslip should be attached using oxygen plasma bonding. Cast chamber is 50 mm x 25 mm x 5 mm. This chamber can be connected to a pump using blunted needles or used as a static chamber using ports sealed with PDMS chamber plugs. Recommended print settings:

- Material: Any PLA or PETG
- Print Temperature: Manufacture's recommended settings
- 0.4 mm nozzle
- 0.05 mm layer thickness
- Ironing activated
- 100% infill

***PDMS_plugs.stl:*** Plugs for sealing the loading ports of all the above PDMS chambers.

- Material: Black food grade PLA
- Print Temperature: Manufacture's recommended settings (typically 215C nozzle, 60C bed)
- 0.4 mm nozzle
- 0.2 mm layer thickness
- 100% infill

Leiden Folder:
This folder contains designs for magnetic 35 mm and 42 mm reusable Leiden chambers. Full print and assembly instructions can be found in the “readme.txt” file in this folder.
